# Effect of irradiation on the survival and susceptibility of female *Anopheles arabiensis* to natural isolates of *Plasmodium falciparum*

**DOI:** 10.1101/2020.01.27.919530

**Authors:** Edwige Guissou, Serge Poda, François de Sales Domombabele Hien, Serge Rakiswende Yerbanga, Dari Frédéric Yannick Da, Anna Cohuet, Florence Fournet, Olivier Roux, Hamidou Maiga, Abdoulaye Diabaté, Jeremie Gilles, Jérémy Bouyer, Anicet G. Ouédraogo, Jean-Baptiste Rayaissé, Thierry Lefèvre, Kounbobr Roch Dabiré

## Abstract

**Background:** The sterile insect technique (SIT) is a vector control strategy relying on the mass release of sterile males into wild vector populations. Current sex separation techniques are not fully efficient and could lead to the release of a small proportion of females. It is therefore important to evaluate the effect of irradiation on the ability of released females to transmit pathogens. This study aimed to assess the effect of irradiation on the survival and competence of *Anopheles arabiensis* females for *Plasmodium falciparum in* laboratory conditions.

**Methods:** Pupae were irradiated at 95 Gy, a sterilizing dose of gamma-rays from Caesium-137 source, and emerging adult females were challenged with one of 14 natural isolates of *P. falciparum*. Seven days post-bloodmeal (dpbm), irradiated and unirradiated-control females were dissected to assess the presence of oocysts. On 14 dpbm, oocyst rupture in mosquito midguts and sporozoite dissemination in head/thoraces were also examined. Two assays were performed to gauge the effect of irradiation on *An. arabiensis* survival. First, the survivorship of irradiated and unirradiated-control mosquitoes exposed to each parasite isolate was monitored. Second, how parasite infection and irradiation interact to influence mosquito lifespan was also assessed by including a group of uninfected unirradiated mosquitoes.

**Results:** Overall, irradiation reduced the proportion of infected mosquitoes but this effect was inconsistent among parasite isolates. Second, there was no significant effect of irradiation on the number of developing oocysts. Third, the proportion of ruptured oocysts at 14 dpbm was higher in irradiated-than in control-unirradiated females, suggesting that irradiation might speed up parasite development. Fourth, irradiation had varying effects on female survival with either a negative effect (assay 1) or no effect (assay 2).

**Conclusion:** Combining these effects into an epidemiological model could help quantifying the net effect of irradiation on malaria transmission in this system. Together, our data indicate that irradiated female *An. arabiensis* could contribute to malaria transmission, and highlight the need for perfect sexing tools which would prevent the release of females as part of SIT programs.

## Introduction

The worldwide annual incidence of malaria declined by 36 % between 2000 and 2015 (Cibulskis *et al.*, 2016). Control measures based on vector management have played an important role in reducing malaria transmission with, for example, the use of long-lasting insecticide-treated nets contributing to an estimated 68 % of the decline in *Plasmodium falciparum* incidence over this period (Bhatt *et al.*, 2015). Since 2015 however, global progress has stalled, and several African countries are currently experiencing an increase in malaria incidence (WHO, 2018). The reasons for these recent increases are unclear but current vector control techniques are showing some limitations. This may include a loss of motivation in tool use (Pulford *et al.*, 2011), and/or vector adaptations such as physiological and behavioral resistance to insecticides (Ranson *et al.*, 2011; Carrasco *et al.*, 2019).

Although improving the use of existing and available tools is essential for malaria control in the near future, there also is an urgent need for the development and implementation of alternative solutions (Feachem *et al.*, 2019). One of them is based on the Sterile Insect Technique (SIT), which aims to control vector populations by releasing sterile males. SIT relies on the massive production of sterile males by irradiation or chemical treatment and introduction into wild insect populations in order to reduce the number of adults in subsequent generations (Knipling, 1955; Knipling *et al.*, 1968; Robinson *et al.*, 2009). With repeated releases, this approach has proven successful in eliminating some agricultural pest species (Dyck, Hendrichs and Robinson, 2005), and has shown promising in suppressing or reducing the density of disease vectors from islands (Vreysen *et al.*, 2000) or from relatively isolated areas such as urban settings (Bellini *et al.*, 2013). More recently, it allowed eliminating two partially isolated populations of *Aedes albopictus* in Guangzhou, China, when used in combination with the incompatible insect technique (Zheng *et al.*, 2019).

In recent years, the joint FAO/IAEA program has spurred renewed interest in the development of SIT approaches for the control of mosquito-borne diseases (Lees *et al.*, 2015; Flores and O’Neill, 2018). With regard to malaria, *Anopheles arabiensis* has focused much of the scientific attention as this species can display localized, narrow range distribution such as river side (Ageep *et al.*, 2009) or urban areas (Dabiré *et al.*, 2014; Azrag and Mohammed, 2018). Accordingly, the radiation biology of this species has been relatively well studied (Helinski, Parker and Knols, 2006; Helinski *et al.*, 2008; Helinski and Knols, 2009; Balestrino *et al.*, 2011; Damiens *et al.*, 2012; Ndo *et al.*, 2014). Besides efficient mass-rearing and optimal level of irradiation ensuring male sterilization with limited impact on sexual competitiveness, a perfect separation technique of male and female mosquitoes prior to release is essential (Mashatola *et al.*, 2018).

To date, the available sexing tools, including pupal size, addition of toxicants to bloodmeal sources, or development of genetic sexing strains, remain imperfect; and a small proportion of females can escape sexing before irradiation (Papathanos *et al.*, 2009; Ndo *et al.*, 2014; Dandalo *et al.*, 2017; Mashatola *et al.*, 2018). These females will be irradiated with the males and can therefore potentially contribute to malaria transmission when released into wild populations. While efforts to find an effective and operational sex separation technique are maintained, it is important to evaluate the effect of irradiation on the ability of female anopheles to transmit *P. falciparum*. Previous work has focused on the influence of irradiation on a large range of traits including sperm production (Helinski and Knols, 2008, 2009; Damiens, Vreysen and Gilles, 2013), male competitiveness (Helinski and Knols, 2008, 2009; Yamada *et al.*, 2014), male and female longevity (Helinski, Parker and Knols, 2006; Dandalo *et al.*, 2017), insemination rate (Poda *et al.*, 2018), oviposition behavior (Poda *et al.*, 2018), fertility and fecundity (Helinski, Parker and Knols, 2006; Poda *et al.*, 2018) but no study has, to our knowledge, characterized the influence of irradiation on the competence of *An. arabiensis* for *P. falciparum*.

Competence, the mosquito ability to ensure parasite development and transmission, is a combined estimate of parasite infectivity and vector susceptibility to infection. It thus encompasses both mosquito resistance mechanisms used to fight the infection and parasite mechanisms used to overcome the vector’s defenses (Lefevre *et al.*, 2018). The molecular and genetic bases of mosquito competence for malaria parasites have been well characterized for a number of mosquito–parasite associations (Molina-Cruz *et al.*, 2015, 2016) and, there also is a great diversity of ways in which biotic and abiotic environmental factors (temperature, mosquito diet, insecticide exposure, microbial gut flora, etc.) can affect mosquito competence (Lefèvre *et al.*, 2013). As any other environmental factors, radiation has also the potential to influence the competence of *Anopheles* vectors for *P. falciparum.* For example, *Aedes aegypti* mosquitoes exposed to a 5000 roentgen dose of X rays-irradiation and infected with a strain of *P. gallinaceum* showed a 2.7 fold reduction in oocyst number compared to unirradiated infected counterparts (Terzian, 1953), thereby suggesting a potential negative effect of irradiation on mosquito competence for malaria parasites (see also Hahn, Haas and Wilcox, 1950; Ward, Bell and Schneider, 1960). In contrast, a study on anopheles mosquitoes found that adult gamma-irradiated *An. quadrimaculatus* displayed increased susceptibility to the nematode *Dirofilaria uniformis* (Duxbury and Sadun, 1963).

The current study aimed to evaluate the effect of a sterilizing dose of gamma-rays from Caesium-137 source on mosquito competence using the parasite *P. falciparum*, responsible for causing the most severe form of human malaria, and the mosquito *An. arabiensis*, a major vector of *P. falciparum* in Africa. Females of *An. arabiensis* were challenged with sympatric field isolates of *P. falciparum* (14 distinct isolates in total) using direct membrane feeding assays and, through a series of experiments, the effects of irradiation on (i) mosquito competence at two distinct time points over the course of infection (oocyst and sporozoite parasite developmental stages), (ii) the timing of oocyst rupture and sporozoite dissemination, and (iii) female survival, were examined.

## Methodology

### Mosquitoes

Laboratory-reared *An. arabiensis* were obtained from an outbred colony established in 2016 and repeatedly replenished with F1 from wild-caught mosquito females collected in Dioulassoba, a central urban area of Bobo-Dioulasso, south-western Burkina Faso, and identified by routine PCR-RFLP (Fanello, Santolamazza and Della Torre, 2002). Mosquitoes were held in 30 × 30 × 30 cm mesh-covered cages under standard insectary conditions (27 ± 2°C, 70 ± 5 % relative humidity, 12:12 LD).

### Irradiation

Irradiation was performed as described in (Poda *et al.*, 2018). Prior to irradiation, pupae were transferred from their rearing trays to plastic cups (Ø = 45mm, h = 85mm) at similar densities. Cups were randomly assigned to one of two treatment groups: irradiation or control-unirradiated. The pupae density in cups was similar between the two treatment groups and did not exceed 200 pupae per cup. One cm of water was left at the bottom of each cup to limit radiation absorbance by water. Pupae were irradiated at a dose of 95.4 ± 0.9 Gy (mean±se) in a Gamma Cell ^137^Cs self-contained gamma source at a rate of 4Gy/min. In *An. arabiensis* males, this dose induces a high level of sterility (Helinski, Parker and Knols, 2006; Poda *et al.*, 2018) while preserving relatively high competitiveness (Helinski, Parker and Knols, 2006). Cups were placed at the center of the irradiation chamber to maximize dose uniformity within the batch. A dosimetry system was used to measure the accurate dose received by each batch using a Gafchromic^®^ HD-V2 film (Ashland, Bridgewater, NJ, USA) placed on the wall of the cups. After irradiation, the optical density of irradiated films was read at both 458nm and 590nm with a dose reader (Dosereader4, Radgen, Budapest, Hungary) and compared to a control. The control group was manipulated in the same way as the irradiated group but was not irradiated. Pupae were then placed in 30 × 30 × 30 cm mesh-covered cages and kept under standard insectary conditions (27 ± 2°C, 70 ± 10 % RH, 12:12 LD) for emergence. Female and male mosquitoes were maintained together on a 5 % glucose solution. Between 3 and 6 days after emergence, control and irradiated females were transferred to cardboard cups (Ø = 75mm, h = 85mm) at a density of 60 mosquitoes per cup.

### Parasite isolates and mosquito experimental infection

Irradiated and control mosquito females were challenged by using blood drawn from naturally *P. falciparum* gametocyte-infected patients recruited among 5–12-year-old school children in villages surrounding Bobo-Dioulasso, Burkina Faso, using Direct Membrane Feeding Assays (DMFA) as previously described (Ouédraogo *et al.*, 2013; Hien *et al.*, 2016). Briefly, thick blood smears were taken from each volunteer, air-dried, Giemsa-stained, and examined by microscopy for the presence of *P. falciparum* at the IRSS lab in Bobo-Dioulasso. Asexual trophozoite parasite stages were counted against 200 leucocytes, while infectious gametocytes stages were counted against 1000 leukocytes. Children with asexual parasitemia of > 1,000 parasites per microliter (estimated based on an average of 8000 leucocytes/ml) were treated in accordance with national guidelines. Asymptomatic *P. falciparum* gametocyte-positive children were recruited for the study.

Gametocyte carrier blood was collected by venipuncture into heparinized tubes. To test for a possible interaction between the natural blocking immunity of the human host (Gouagna *et al.*, 2004; Da *et al.*, 2015; Stone *et al.*, 2018) and the irradiation on mosquito infection, DMFA were performed using either whole donor blood or with replacement of the serum by a non-immune AB serum (see Additional file 1). Mosquitoes were starved of glucose solution for 12 h prior to the exposure. Three to six day old female mosquitoes, emerged from irradiated or control pupae, were allowed to feed on this blood for one hour. Non-fed or partially fed females were removed and discarded, while the remaining fully-engorged mosquitoes were maintained in a biosafety room under standard insectary conditions (27 ± 2°C, 70 ± 10 % RH, 12:12 LD). Mosquitoes were provided with a sugar meal consisting in a 5 % glucose solution on cotton wool following blood-feeding.

### Experiment 1: Effects of irradiation on *An. arabiensis* competence for *P. falciparum*

Competence was characterized by infection prevalence (i.e. the proportion of mosquitoes that develop infection upon feeding on an infectious bloodmeal) and intensity (i.e. the average number of parasites among infected mosquitoes). Infection prevalence and intensity were here gauged at two distinct points in time over the course of infection (Table 1):

i. On day 7 post-bloodmeal (dpbm), the midguts of a total of 383 irradiated females and 378 control females fed with blood from one of 8 gametocyte carriers (Table 1) were dissected, stained with 2 % mercurochrome, and the presence and number of oocysts (immature, non-transmissible stage of malaria parasites) were recorded using light under the microscopy (×400).
ii. On 14 dpbm, the heads and thoraces of a total of 473 irradiated and 489 control females fed with blood from one of 10 gametocyte carriers (Table 1) were dissected, and the presence and quantity of sporozoites (mature transmissible stage) were determined using qPCR (see below).

**Table 1:** Summary description of the experiments.

| Experiment | Time point | Parasites isolates (gam/µl) | Measured traits | Total sample size (N total)<br>Mean ± se (median) [range]<br>number of mosquitoes for each parasite isolate |  |
| --- | --- | --- | --- | --- | --- |
|  |  |  |  | irradiated | unirradiated-control |
| **Experiment 1:**<br>Effects of irradiation on *An. arabiensis* competence for *P. falciparum* | 7 dpbm | A (64), C (160), D (88), E (32), G (48), H (56), I (48), J (32) | **Oocyst prevalence:** the number of mosquitoes harboring at least one oocyst in their midguts out of the total number of dissected mosquitoes | N total = 383<br>47.875 ± 4.9 (50) [21-71] | N total = 378<br>47.25 ± 4.9 (51) [23-66] |
|  |  |  | **Oocyst intensity:** the mean number of oocysts in infected mosquitoes | N total = 180<br>19 ± 2.7 (19.5) [5-31] | N total = 152<br>22.5 ± 3.22 (20) [13-37] |
|  | 14 dpbm | E (32), H (56), I (48), J (32), K (72), L (168), M (32), N (136), O (96), P(96) | **Sporozoite prevalence:** the number of mosquito head/thorax detected positive to *P. falciparum* using qPCR out of the total number of dissected head/thoraces | N total = 473<br>47.3 ± 3.7 (50) [17-55] | N total = 489<br>48.9 ± 4.35 (48.5) [27-78] |
|  |  |  | **Sporozoite intensity:** The mean number of amplification cycle during qPCR (the lower the Ct, the higher the sporozoite intensity) | N total = 257<br>25.7 ± 2.7 (26) [9-38] | N total = 248<br>24.8 ± 4.7 (23.5) [7-59] |
| **Experiment 2:**<br>Effects of irradiation on *P. falciparum* oocyst rupture in mosquito guts and sporozoite dissemination in head/thoraces | 14 dpbm | K (72), L (168), M (32), N (136), O (96), P (96) | **Proportion of infected mosquitoes with ruptured oocysts:** the number of mosquitoes with at least one ruptured oocyst out of the total number of infected mosquitoes (i.e. harboring either intact and/or ruptured oocysts) | N total = 276<br>24 ± 5 (23) [12-44] | N total = 243<br>20.7 ± 4 (18.5) [9-37] |
|  |  |  | **Proportion of ruptured oocysts:** the number of ruptured oocysts out of the total number of oocysts (intact + ruptured) |  |  |
|  |  |  | **Proportion of oocyst-infected mosquitoes with sporozoites in their head and thorax:** the number of oocyst-infected mosquitoes harboring sporozoites in their head/thoraces out of the total number of infected mosquitoes |  |  |
| **Experiment 3:**<br>Effects of irradiation on *An. arabiensis* survival | 1-7 dpbm | A (64), C (160), D (88), G (48) | From 1 to 7 dpbm, the number and time of death was recorded among mosquitoes exposed to the infectious blood-meal | N total = 165<br>41.25 ± 11 (35.5) [22-72] | N total = 168<br>42 ± 8.7 (41.5) [24-61] |
|  | 1-14 dpbm | E (32), H (56), I (48), J (32), K (72), L (168), M (32), N (136), O (96), P(96) | From 1 to 14 dpbm, the number and time of death was recorded among mosquitoes exposed to the infectious blood-meal | N total = 880<br>88 ± 9.9 (85.5) [31-146] | N total = 803<br>80.3 ± 11.9 (62.5) [41-137] |
|  | 1-35 dpbm | J (32) | From 1 dpbm until all mosquitoes had died (35 dpbm), the number and time of death was recorded among both infected mosquitoes and uninfected control mosquitoes | infected = 26<br>uninfected = 49 | infected = 14<br>uninfected = 45 |

### Experiment 2: Effects of irradiation on *P. falciparum* oocyst rupture in mosquito midguts and sporozoite dissemination in head/thoraces

On 14 dpbm, 276 irradiated and 243 control mosquito females fed an infectious blood from one of 6 gametocyte carriers were dissected for the microscopic observation of oocysts in midguts and the qPCR detection of sporozoites in head/thoraces (Table 1). Oocyst rupture in mosquito midgut and sporozoite invasion of salivary glands is highly asynchronous: while some oocysts are intact and keep developing on 14 dpbm, others have already ruptured and released their sporozoites. To explore possible difference in the timing of sporozoite dissemination in mosquito salivary glands between irradiated and control females, three traits were measured:

i. the proportion of infected mosquitoes with ruptured oocysts on 14 dpbm. This is the number of mosquitoes with at least one ruptured oocyst in their midguts at 14 dpbm out of the total number of infected mosquitoes (i.e. harboring either intact and/or ruptured oocysts);
ii. the proportion of ruptured oocysts at 14 dpbm. This is, for each infected mosquito, the number of ruptured oocysts out of the total number of oocysts (intact + ruptured);
iii. the proportion of oocyst-infected mosquitoes with sporozoites in their head and thorax at 14 dpbm. This is the number of oocyst-infected mosquitoes harboring sporozoites in their head/thoraces at 14 dpbm out of the total number of infected mosquitoes (i.e. harboring either intact and/or ruptured oocysts).

### Experiment 3: Effects of irradiation on *An. arabiensis* survival

Two assays were performed to gauge the effect of irradiation on *An. arabiensis* survival. First, as part of the previous experiments, the survivorship of irradiated and unirradiated-control mosquitoes exposed to each parasite isolate (n = 14 isolates) was monitored from 1 to 7 days post-treatment (isolates A, C, D and G) or from 1 to 14 dpbm (isolates E, H, I, J, K, L, M, N, O, P). Every morning at 08:00, dead mosquitoes were removed and counted from each cage. The remaining alive mosquitoes used for midgut dissection at 7 and/or 14 dpbm (experiment 1) were considered in the analysis and given a censoring indicator of “0”.

Second, to determine how parasite infection and irradiation interact to influence mosquito longevity, a membrane feeding assay was performed following the same general procedure as described above except that a group of uninfected control mosquitoes was added, and that survival was monitored until all the mosquitoes had died. Uninfected control mosquitoes received heat-treated gametocytic blood to kill the parasite gametocytes as previously described (Sangare *et al.*, 2013; Hien *et al.*, 2016; Nguyen *et al.*, 2017). For each group (irradiated-parasite exposed, irradiated-parasite unexposed, control-parasite exposed and control-parasite unexposed), between 40 and 60 females were placed in one of two 20 × 20 × 20 cm cages to avoid possible cage effect on mosquito survival. Females were fed a 2.5 % glucose solution every other day and provided water-soaked cottons ad libitum. Dead mosquitoes were counted from each cage (n = 8 cages) every morning at 8:00 and individually stored at −20°C to determine their infection status using qPCR (see below).

### *Plasmodium falciparum* DNA extraction and qPCR

*P. falciparum* genomic DNA was extracted from head-thorax mosquitoes by grinding tissues with a micro pestle in an extraction buffer (0.1 M Tris HCl, pH 8.0, 0.01 M EDTA, 1.4 M NaCl, 2 % cetylltrimethyl ammonium bromide). The mixture was incubated at 65°C for ten min. Total DNA was extracted with chloroform, precipitated in isopropanol, washed in 70 % ethanol, and resuspended in sterile water (Morlais *et al.*, 2004). Parasite detection was carried out by qPCR, using *P. falciparum* mitochondrial DNA specific primers 5’-TTACATCAGGAATGTTTTGC-3’ and qPCR-PfR 5’-ATATTGGGATCTCCTGCAAAT-3’ (Boissière *et al.*, 2013).

### Statistical analyses

All statistical analyses were performed in R (version 3.6.1). Logistic regression by generalized mixed linear models (GLMM, binomial errors, logit link; lme4 package) were used to test the effect of irradiation on (i) the prevalence of oocysts and sporozoites (experiment 1), (ii) the proportion of infected mosquitoes with ruptured oocysts (experiment 2), (iii) the proportion of ruptured oocysts (experiment 2), (iv) the proportion of oocyst-infected mosquitoes with sporozoites in their head and thorax (experiment 2). A GLMM with zero truncated negative binomial errors (glmmTMB package) was used to test the effect of irradiation on the oocyst intensity (experiment 1). A GLMM with Gaussian distribution (lme4 package) was used to test the effect of irradiation on the sporozoite intensity (Ct: mean number of amplification cycle during qPCR, experiment 1). For each GLMM, the full model included irradiation treatment (irradiated vs. unirradiated-control) and gametocytemia (the mean number of gametocytes in parasite isolates) as fixed effects and parasite isolate as a random effect. The effect of irradiation on mosquito survivorship (survival assay 1) was analyzed using a Cox’s proportional hazard regression mixed model (coxme package) with censoring and with parasite isolate set as a random factor. The effect of irradiation and infection on mosquito survivorship (survival assay 2) was analyzed using Cox’s proportional hazard regression mixed model without censoring and with cage identity set as a random factor. Model simplification used stepwise removal of terms, followed by likelihood ratio tests (LRT). Term removals that significantly reduced explanatory power (P < 0.05) were retained in the minimal adequate model.

### Ethical considerations

The selection of parasite isolate was made from asymptomatic gametocyte carriers recruited among 5-12 year old children in the villages of the medical district of Dandé and Soumousso according to the protocol approved respectively by the Centre Muraz and IRSS ethics committees: A003-2012/CE-CM and 2017-003/MESRSI/CNRST/IRSS/CEIRES. Prior to inclusion, informed consent was obtained from parents or legal guardian. The protocol was in line with the 2002 Helsinki Declaration on Ethical Principles for Medical Research Involving Human Subjects.

## Results

### Experiment 1: Effects of irradiation on *An. arabiensis* competence for *P. falciparum* Oocyst prevalence and intensity at day 7 post-bloodmeal

Irradiation reduced the proportion of infected mosquitoes by 16.8 % (180 infected control mosquitoes/378 = 47.6%; and 152 infected irradiated mosquitoes / 383 = 39.6%; *LRT X^2^_1_* = 5.2; P = 0.02; Figure 1A). Although no significant effect of gametocytemia on oocyst prevalence was found (*LRT X^2^_1_* = 0.2; P = 0.65), there was an interaction between irradiation and gametocytemia (*LRT X^2^_1_* = 19.5; P<0.001). In particular, while irradiation reduced mosquito infection rate of parasite isolates C, D, G, I, it had no effect on A, E and even slightly increased the infection rate of isolates H and J (Figure 1A).

**Figure 1:**
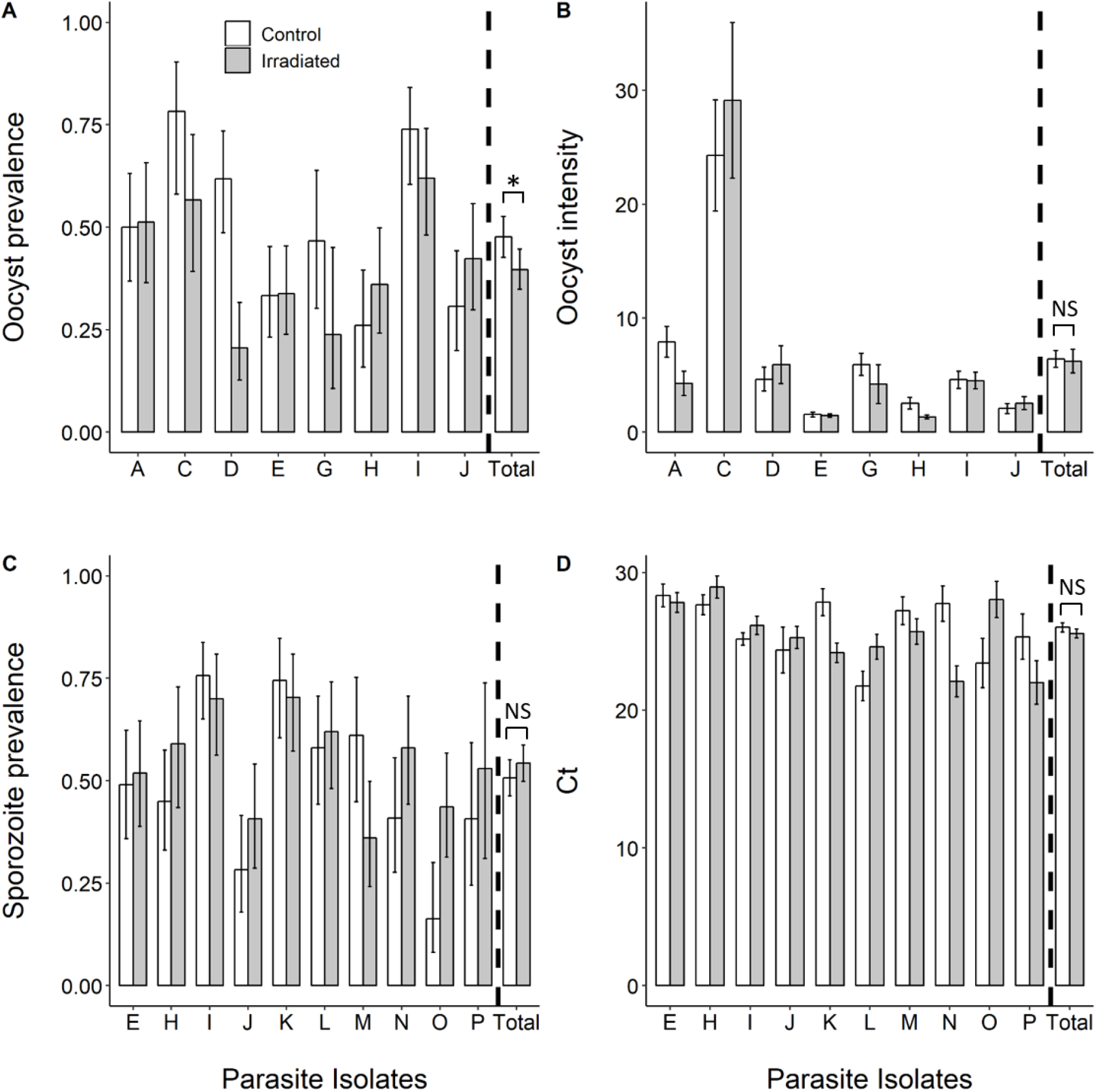
Effect of irradiation on the competence of *Anopheles arabiensis* for natural isolates of *P. falciparum*. (A) Oocyst prevalence (± 95 % CI) on day 7 post-bloodmeal (dpbm), expressed as the number of mosquito females harboring at least one oocyst in their midguts out of the total number of dissected females, for each treatment (white bars: unirradiated-control mosquitoes, grey bars: irradiated mosquitoes) and for 8 parasite isolates. (B) Infection intensity (± se) at 7 dpbm, expressed as the mean number of developing oocysts in the guts of infected females, for each treatment and 8 parasite isolates. (C) Sporozoite prevalence (± 95 % CI) at 14 dpbm, expressed as the number of mosquito head/thoraces detected positive to *Plasmodium falciparum* using qPCR out of the total number of dissected head/thoraces, for each treatment and for 10 parasite isolates. (D) Sporozoite intensity at 14 dpbm, expressed as the mean number (± se) of amplification cycle during qPCR (the lower the Ct, the higher the sporozoite intensity) for each treatment and for 10 parasite isolates. * denotes statistically significant difference (P value: 0.01< * < 0.05); NS: not significant.

The mean number of developing oocysts in infected females (i.e. intensity) was not significantly affected by irradiation (*LRT X^2^_1_* = 0.0017; P = 0.97, Figure 1B). Gametocytemia had no effect on intensity (*LRT X^2^_1_* = 0.54; P = 0.46, Figure 1B). There was a significant interaction between gametocytemia and treatment (LRT X_2_^1^ = 9.58, P = 0.002, Figure 1B) such that irradiation either decreased (isolates A, G, H), increased (C, D) or had no effect (E, I, J) on oocyst intensity.

### Sporozoite prevalence and intensity at day 14 post-bloodmeal

The proportion of mosquitoes with disseminated sporozoites in their head/thorax was similar between irradiated and control females (control: 248/489 = 50.7 ± 4 %; irradiated: 257/473 = 54.3 ± 5 %, *LRT X^2^_1_* = 2.56, P = 0.11; Figure 1C). There was no effect of gametocytemia on sporozoite prevalence (*LRT X^2^_1_* = 0.12, P = 0.73, Figure 1C), and a marginally non-significant interaction between irradiation and gametocytemia (*LRT X^2^_1_* = 3.5, P = 0.06, Figure 1C).

The mean number of amplification cycle during qPCR (the lower the Ct, the higher the sporozoite intensity) did not vary with irradiation (mean Ct irradiated = 25.57 ± 0.32 (n = 257), mean Ct control = 26.02 ± 0.33 (n = 248), *LRT X^2^_1_* = 0.55, P = 0.46, Figure 1D). Gametocytemia had a significant effect on sporozoite intensity (*LRT X^2^_1_* = 7.7, P = 0.006), with higher gametocyte density in blood leading to an increase in sporozoite density in mosquito head and thoraces. Finally, there was no interaction between irradiation and gametocytemia on sporozoite intensity (*LRT X^2^_1_* = 0.04, P = 0.85).

### Experiment 2: Effects of irradiation on *P. falciparum* oocyst rupture in mosquito guts and sporozoite dissemination in head/thoraces

Uninfected mosquitoes were excluded from the analysis and the parasite oocyst rupture in mosquito guts and sporozoite dissemination to head/thoraces were compared between irradiated and control infected individuals (N irradiated = 144/276 (52 %), N control = 124/243 (51 %)). Among these infected mosquitoes, the proportion of individuals with at least one ruptured oocyst in their midgut at 14 dpbm was higher in irradiated females than in control counterparts (*LRT X^2^_1_* = 5.8, P = 0.016, Figure 2A). In particular, 86 % (124/144) of irradiated infected mosquitoes had at least one ruptured oocyst in their midguts, while only 75 % (93/124) of control infected females exhibited ruptured oocysts. This result suggests that the release of sporozoites from oocysts happened earlier in irradiated than in control females.

**Figure 2:**
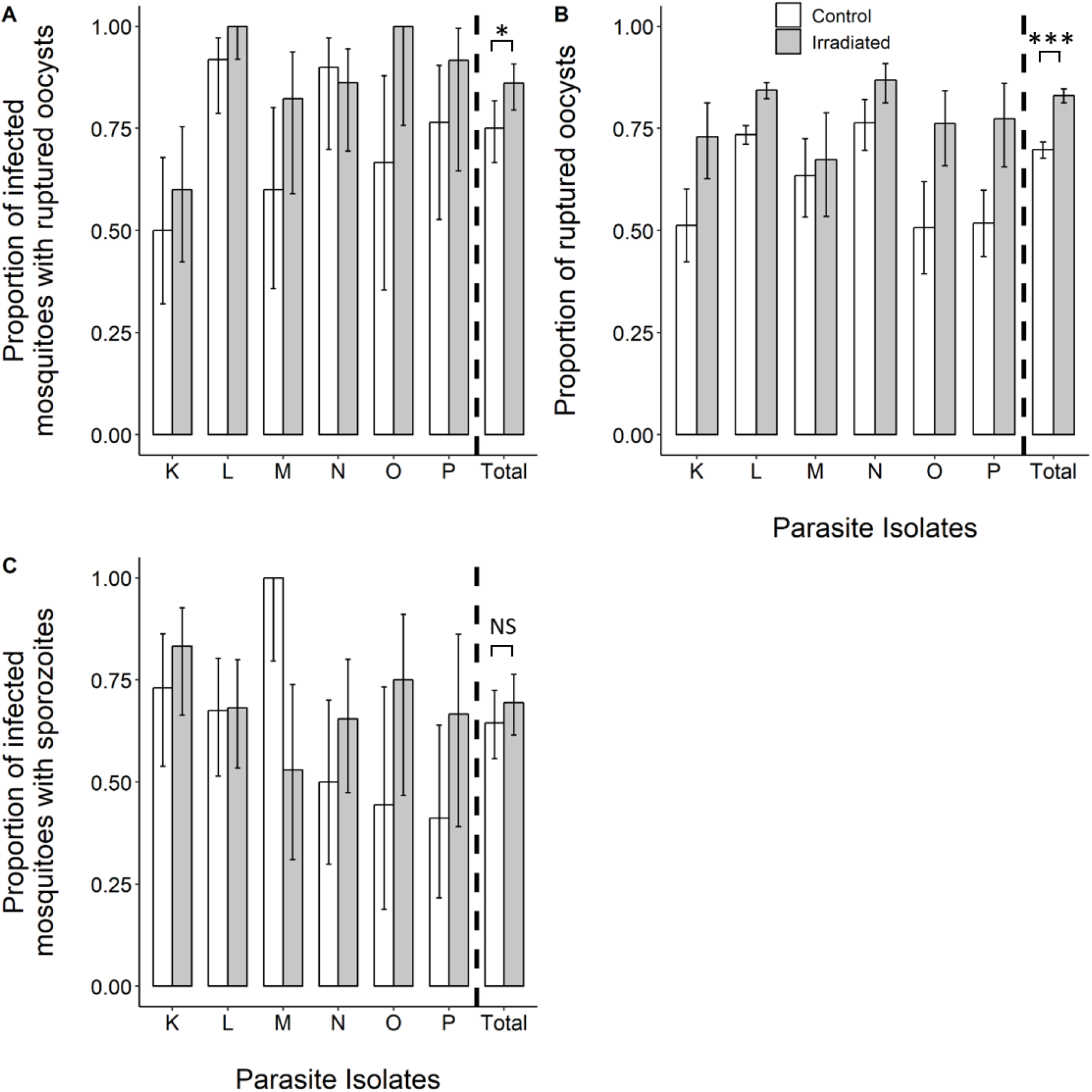
Effect of irradiation on *P. falciparum* oocyst rupture in mosquito guts and sporozoite dissemination in head/thoraces on day 14 post-bloodmeal (dpbm). (A) Proportion of infected mosquitoes with ruptured oocysts (± 95 % CI), expressed as the number of mosquitoes with at least one ruptured oocyst out of the total number of infected mosquitoes (i.e. harboring either intact and/or ruptured oocysts) at 14 dpbm for each treatment (white bars: unirradiated-control mosquitoes, grey bars: irradiated mosquitoes) and for 6 parasite isolates. (B) Proportion of ruptured oocysts (± 95% CI), expressed as the number of ruptured oocysts out of the total number of oocysts (intact + ruptured) at 14 dpbm for each treatment and 6 parasite isolates. (C) Proportion of oocyst-infected mosquitoes with sporozoites in their head and thorax (± 95% CI), expressed as the number of oocyst-infected mosquitoes harboring sporozoites in their head/thoraces out of the total number of infected mosquitoes at 14 dpbm, for each treatment and for 6 parasite isolates. * denotes statistically significant difference (P value: 0.01< * < 0.05; 0 < *** < 0.001); NS: not significant.

In addition, the proportion of ruptured oocysts was higher in irradiated mosquitoes (irradiated: 1509 ruptured oocysts out of a total of 1817 counted oocysts (83 %), controls: 1443 ruptured oocysts out of a total of 2068 counted oocysts (69.8 %), *LRT X^2^_1_* = 85, P < 0.001, Figure 2B), further suggesting that irradiation speeded up oocyst maturation and sporozoite release.

Finally, the proportion of oocyst-infected mosquitoes with disseminated sporozoites in their head/thorax was not affected by irradiation treatment (*LRT X^2^_1_* = 2, P = 0.12, Figure 2C). There was no main effect of gametocytemia on the proportion of oocyst-infected mosquitoes with disseminated sporozoites in their head/thorax (*LRT X^2^_1_* = 1.65, P = 0.2). There was a significant interaction between gametocytemia and treatment (*LRT X^2^_1_* = 4.6, P = 0.03), with irradiation either decreasing (isolates M), or increasing (K, N, O, P) the proportion of oocyst-infected mosquitoes with disseminated sporozoites.

### Experiment 3: Effects of irradiation on *An. arabiensis* survival

In the first assay, the survival of females exposed to one of 14 parasite isolates was monitored from 1 to 7 dpbm or from 1 to 14 dpbm (Table1). The overall survival rate from 1 to 7 dpbm (isolates A, C, D, G) was very high, with only 3.9 % of mosquitoes (13 / 333) that died between 1 to 7 dpbm, and there was no survival difference between irradiated and control non-irradiated mosquitoes (LRT *X^2^_1_* = 1, P = 0.31, Figure 3A). However, from 1 to 14 dpbm (isolates E, H, I, J, K, L, M, N, O, P), irradiated mosquitoes died at a higher rate than control mosquitoes (mortality rate irradiated: 21.25 % (187 / 880), control: 11.71 % (94/803), LRT *X^2^_1_* = 22.3, P < 0.001, Figure 3B).

**Figure 3:**
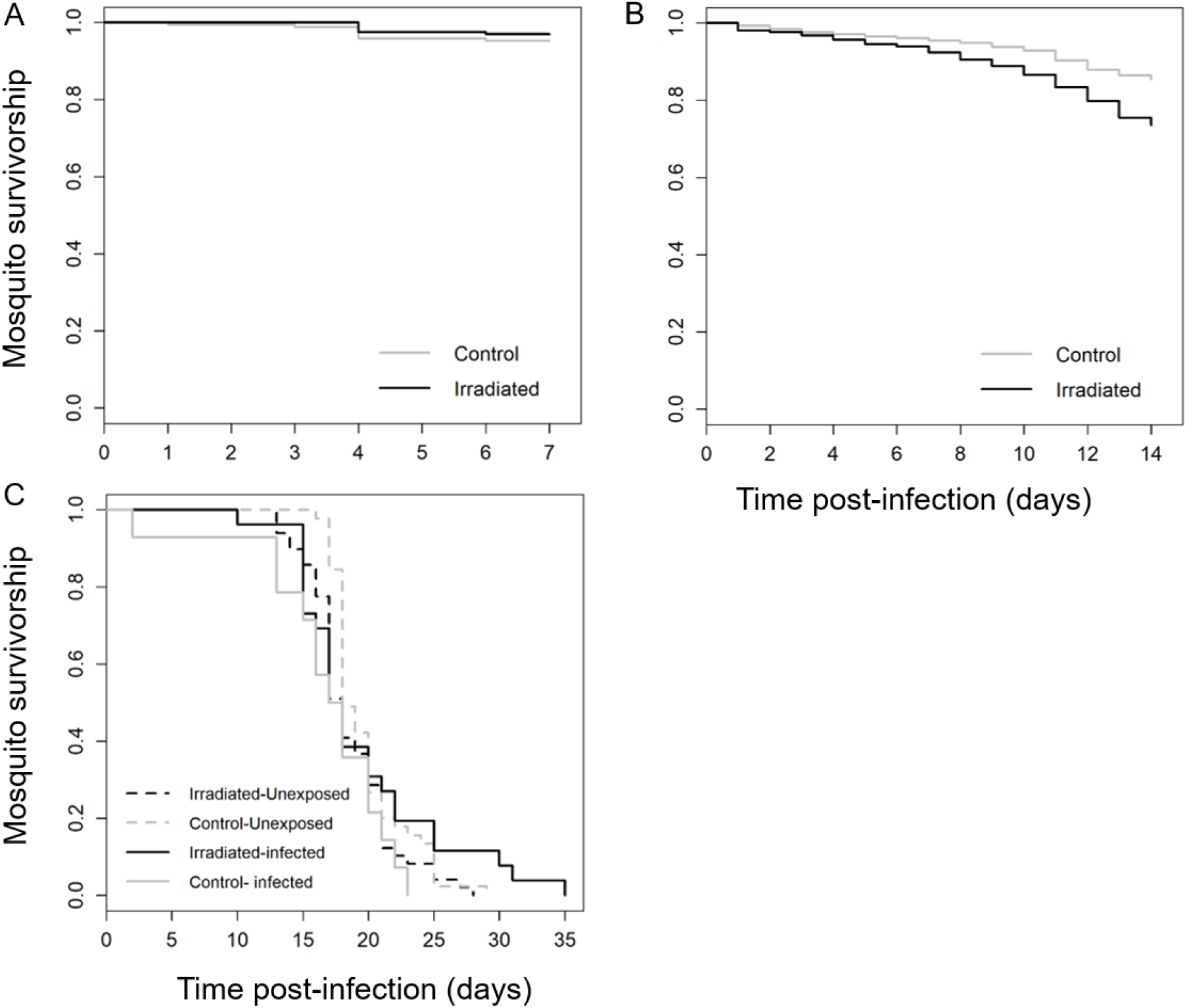
Effect of irradiation on the survival of *Anopheles arabiensis*. (A) Survivorship of malaria-exposed mosquitoes from 1 to 7 dpbm for each treatment (grey line: unirradiated-control, black line: irradiated) using 4 parasite isolates. (B) Survivorship of malaria-infected mosquitoes from 1 to 14 dpbm for each treatment using 10 parasite isolates. (C). Survivorship of both malaria-infected (solid lines) and uninfected unirradiated (dahes lines) mosquitoes from 1 to 35 dpbm for each treatment (grey: unirradiated-control, black: irradiated) using 1 parasite isolate.

In the second assay, the survival of irradiated mosquitoes exposed to parasites (n = 55), irradiated unexposed (n = 49), unirradiated exposed (n = 52) and unirradiated unexposed (n = 45) females was monitored from 1 to 35 dpbm, when the last mosquito died. The DNA of parasite-exposed dead mosquitoes was extracted to detect the presence of *P. falciparum* using qPCR. Mosquitoes (irradiated or non-irradiated) which remained uninfected upon parasite exposure were excluded from the analysis to focus on the effect of infection and irradiation on mosquito survival. In this second assay using smaller number of mosquitoes (Table 1), there was no effect of irradiation on mosquito survival (LRT *X^2^_1_* = 0.04, P = 0.84, Figure 3C). Infection did not significantly reduce mosquito survival (LRT *X^2^_1_* = 0.05, P = 0.82, Figure 3C). Finally, there was a marginally significant interaction between irradiation and infection (LRT *X^2^_1_* = 4, P = 0.045, Figure 3C), such that irradiation resulted in an increased lifespan in infected mosquitoes but caused a reduced lifespan in uninfected mosquitoes.

## Discussion

Our data shows that irradiation had contrasting effects on critical parameters affecting the capacity of *An. arabiensis* to transmit *P. falciparum*, including mosquito competence, the parasite development time and survival. First, irradiation reduced the proportion of mosquitoes harboring parasite oocysts upon ingestion of bloodmeals from gametocyte carriers. Second, irradiation increased both the proportion of mosquitoes with ruptured oocysts and the proportion of ruptured oocysts in mosquito guts at 14 dpbm. Third, irradiation either decreased (survival assay 1) or had no effect (assay 2) on the lifespan of *An. arabiensis* females. While reduced mosquito competence and survival would limit *An. arabiensis* vectorial capacity, shorter parasite development time would tend to increase it. Combining these effects into an epidemiological model could help quantifying the net effect of irradiation on malaria transmission in this system.

Although irradiated females displayed reduced oocyst infection rate compared to non-irradiated individuals, the parasite development was not fully suppressed. If released into the wild, irradiated females will therefore likely contribute to malaria transmission, provided that irradiation does not impair the host-seeking and -feeding behaviors of these females. Our results therefore highlight the need for perfect sexing tools which would prevent the release of females as part of SIT programs.

The precise mechanisms behind irradiation-mediated reduction of *Plasmodium* infection are not yet clear but interferences with mosquito immunity, microbiota and/or parasite infectivity mechanisms are likely. Although it is well-known that irradiation causes DNA damages, oxidative stress, and changes in gene expression including immune genes (Zhikrevetskaya *et al.*, 2015), its impact on insect host-pathogen interactions remain generally unclear (Morley, 2012). While a study found that irradiated *Tephritidae* flies displayed damaged midgut and peritrophic membrane resulting in decreased bacterial growth (Lauzon and Potter, 2012), irradiated *Spodoptera* butterflies showed increased susceptibility to a nucleopolyhedrosis virus (Sayed and El-Helaly, 2018). Similarly, in mosquito-malaria parasites associations, X-ray irradiation caused increased *Ae. aegypti* resistance to *P. gallinaceum* (Terzian, 1953; Ward, Bell and Schneider, 1960), while gamma-ray irradiation enhanced the development of *Dirofilaria uniformis* in *Anopheles quadrimaculatus* (Duxbury and Sadun, 1963). Together, the few existing studies on this topic suggest that the observed changes in infection level are mediated mostly through radiation damage to the insect midgut rather than through altered immune response such as hemocyte production (Jafri, 1965; Christensen, Huff and Li, 1990; Morley, 2012). In addition, the effects of irradiation on infection seem to be dose-dependent. For example, at a dose of 1000 r of x-ray, the competence of *Ae. aegypti to P. gallinaceum* decreased by only 1.15 times compared to unirradiated-control mosquitoes; while at doses between 5,000 and 40,000 r, competence decreased by a factor of 2.75 to 4 (Terzian, 1953). Further investigations are required to determine whether the decreased susceptibility of irradiated *An. arabiensis* to *P. falciparum* oocysts is also dose-dependent.

In this study, the effect of irradiation on mosquito infection strongly varied among parasite isolates (Figure 1). Why irradiation reduced *An. arabiensis* competence for some parasite isolates and not others is unclear. We first postulated that the natural blocking immunity of the human host could play a role. To test this possibility, the natural serum of isolates K to P was replaced by naive AB serum (Gouagna *et al.*, 2004; Da *et al.*, 2015; Stone *et al.*, 2018) (Additional file 1). Similar to assays using unchanged natural serum (isolates A to J), assays with serum replacement showed either increased (L, N, O, and P) or decreased (K and M) infection in irradiated mosquitoes (Additional file 1: Figure S2). Because the characterization of vector competence for oocyst and sporozoite stages partly relied on different gametocyte carriers (Table 1), such isolate-dependent effect of irradiation could also explain why, on average, the sporozoite infection rate of irradiated individuals was not significantly lower than that of unirradiated-control individuals (Figure 1C). Here, we used wild parasite isolates from a geographic area characterized by an important genetic diversity (Somé *et al.*, 2018). Accordingly, some parasite clones might perform well in irradiated mosquitoes while others would be more infective to non-irradiated mosquitoes. Future genotyping studies of the parasite population used to perform the experimental infections of irradiated mosquitoes would be required to explore this possibility.

Our results suggest an earlier sporozoite invasion of salivary glands among irradiated females. This is supported by the higher proportion of infected mosquitoes with ruptured oocysts (Figure 2A), the higher proportion of ruptured oocysts (Figure 2B), and the higher proportion (although not significant) of infected mosquitoes with sporozoites at 14 dpbm (Figure 2C). Gamma-irradiation might speed up *Plasmodium* development within anopheles vectors. Shorter parasite’s Extrinsic Incubation Period (EIP) following insect host irradiation was previously described in *Trypanosoma spp* – infected tsetse flies (Moloo, 1982). In this system, the parasite migration to the haemocoel occurred earlier in irradiated than in unirradiated-control flies, possibly because of changes in the ultrastructural organization of the insect gut (Stiles *et al.*, 1989). Exploring the temporal dynamics of *P. falciparum* development using mosquitoes dissected at different time points during the course of infection would provide more detailed and robust information. The number of mosquitoes in our experiments was insufficient to perform such temporal monitoring of the EIP and future experiments are required to confirm our observations made at 14 dpbm.

The effects of irradiation on the survival of *An. arabiensis* females were inconsistent. In our first assay, the monitoring of 165 irradiated and 168 unirradiated-control females from 1 to 7 dpbm following the ingestion of a gametocyte-infected bloodmeal revealed no effect of irradiation. Within this period, mosquito survival was very high with only 8 deaths in the unirradiated-control group and 5 in the irradiated group. However, when the monitoring expanded to 14 dpbm on much bigger sample size (880 irradiated and 803 unirradiated-control females), the irradiated group recorded twice as many deaths as the unirradiated-control group (21.25 % vs 11.71 %). Finally, no significant influence of irradiation was observed as part of the second survival assay in which 26 infected-irradiated, 49 uninfected-irradiated, 14 infected-unirradiated and 45 uninfected-unirradiated mosquitoes were monitored until all individuals had died. Unlike the first assay in which mosquitoes were maintained on a 5 % glucose solution ad libitum, mosquitoes received a 2.5 % glucose solution every other day in this second assay. This was supposed to induce nutritional stress in mosquitoes and help to better detect possible effects of radiation on survival (Roux *et al.*, 2015; Poda *et al.*, 2018). Inconsistent effects of irradiation on the survival of mosquito females were previously observed, with some studies reporting either lifespan reduction (Terzian, 1953; Brelsfoard, St Clair and Dobson, 2009), no effect (Darrow, 1968; Wakid *et al.*, 1976; Brelsfoard, St Clair and Dobson, 2009; Dandalo *et al.*, 2017) or even increase (Brelsfoard, St Clair and Dobson, 2009). For example, in the mosquito *Ae. polynesiensis*, irradiation of females <24 hrs post-pupation at 20 Gy and 40 Gy induced a significant lifespan reduction compared to non-irradiated females, while irradiation at 30 Gy had no effect and irradiation at 40 Gy of females > 24 hrs post-pupation boosted female lifespan. If confirmed in field conditions, the irradiation-mediated reduction of mosquito lifespan observed from 1 to 14 dpbm would not be strong enough to prevent the completion of *Plasmodium* incubation period and hence the contribution of these females to malaria transmission (Brelsfoard, St Clair and Dobson, 2009).

## Conclusion

Our data indicate that irradiation of female *An. arabiensis* can reduce competence and survival, but not to the point of preventing malaria transmission. Irradiated females therefore remain potential vectors and further studies are required to develop fully effective sexing tools to prevent possible releases of irradiated females into the wild. Until we find such sexing tools, it will be important to expand our knowledge on the radiation biology of female mosquito vectors.

## Abbreviations

dpbm: days post-bloodmeal
SIT: Sterile Insect Technique
L:D: Light Dark
qPCR: Real-time Polymerase Chain Reaction
DMFA: Direct Membrane Feeding Assay

## Acknowledgements

We would like to thank all volunteers for participating in this study as well as the local authorities for their support. We are very grateful to the IRSS staff in Burkina Faso for technical assistance.

## Availability of data and materials

The raw datasets are available from the corresponding author

## Authors’ contributions

EG, TL, KRD conceived and designed the study. EG and TL drafted the manuscript. EG and TL analysed the data. EG, SP, FdSDH conducted the experiments. JBR provided access to irradiation facilities. OR, TL, JG, JB and KRD supervised the study. All authors read, revised and approved the final manuscript.

## Ethics approval and consent to participate

The protocol was approved by the Centre Muraz and IRSS ethics committees: A003-2012/CE-CM and 2017-003/MESRSI/CNRST/IRSS/CEIRES. Prior to inclusion, informed consent was obtained from the parents or legal guardian of the volunteers.

## Consent for publication

NA

