## Additional file 1 for "Effect of irradiation on the survival and susceptibility of female *Anopheles arabiensis* to natural isolates of *Plasmodium falciparum*"

Infection of human hosts with *Plasmodium falciparum* lead to the natural production of antibodies that can reduce parasite infectivity in the mosquito vectors. The concentration of antibody and the resulting natural transmission-blocking immunity is uneven among gametocyte carriers resulting in variable levels of infection in the mosquito (Gouagna *et al.*, 2004; Da *et al.*, 2015; Stone *et al.*, 2018).

In our first set of experiments, we noticed that the effect of irradiation on mosquito infection strongly varied among parasite isolates (Figure 1). We therefore postulated that the natural blocking immunity of the human host could play a role. To test this possibility, the natural plasma of some isolates was replaced by naive AB serum following the methodology described in Da et al. 2015. Such replacement usually increases the infection prevalence and intensity in mosquitoes (Bousema *et al.*, 2011; Da *et al.*, 2015). As part of experiment 1 (see main text), the heads and thoraces of a total of 473 irradiated and 489 control females fed with blood from one of 10 gametocyte carriers (Table 1) were dissected, and the presence and quantity of sporozoites were determined using qPCR. Of these 10 parasite isolates, 4 had unchanged autologous plasma (isolates E, H, I, J) and 6 had replaced naïve AB control serum (K, L, M, N, O, and P). Isolates E and J both had a gametocytemia of 32 gametocytes/ul of blood, H = 56, I = 48. Isolates K showed a gametocytemia of 72 gametocytes/ul of blood, L=168, M=32, N=136, and isolates O and P=96.

Our results show that there was no significant positive relationship between gametocytemia and infection prevalence or intensity (prevalence: $LRT\ X^2_1 = 0.35$, P = 0.55, Figure S1A; intensity: $LRT\ X^2_1 = 3$, P = 0.08, FigS1B) and no significant interaction with the type of serum (prevalence: $LRT\ X^2_1 = 1.2$, P = 0.28, FigS1A; intensity: $LRT\ X^2_1 = 0.002$, P = 0.97, FigS1B). Why higher gametocytemiae do not produce higher levels of infection may first seem unexpected (Da et al. 2015). However, previous work showed that the lack of significant positive correlation is usually observed for gametocytemia < 200 (Churcher *et al.*, 2013).

Second, similar to assays using unchanged natural serum (isolates A to J), assays with serum replacement showed either increased (L, N, O, P) or decreased (K, M) infection in irradiated mosquitoes. This suggests that natural transmission-blocking immunity is not a likely explanation for the observed inconstancies in the effect of irradiation on mosquito infection.

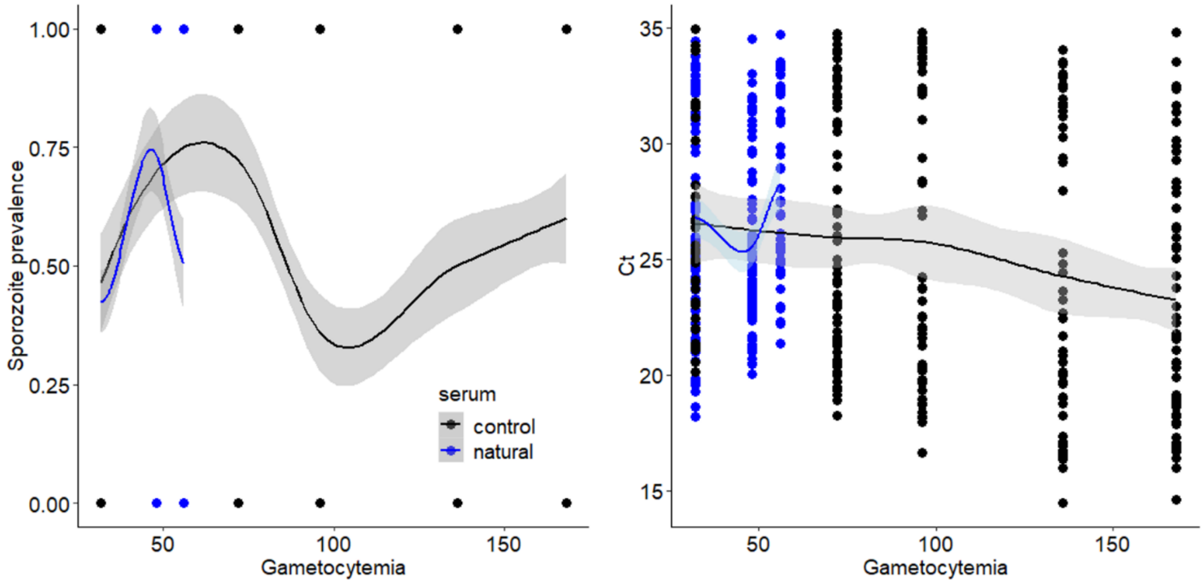


Figure S1: Relationship between Sporozoite prevalence and gametocytemia (left) and between Ct and gametocytemia (right). Blue lines indicates isolates for which the plasma was kept unchanged and black lines isolates with naïve AB serum.

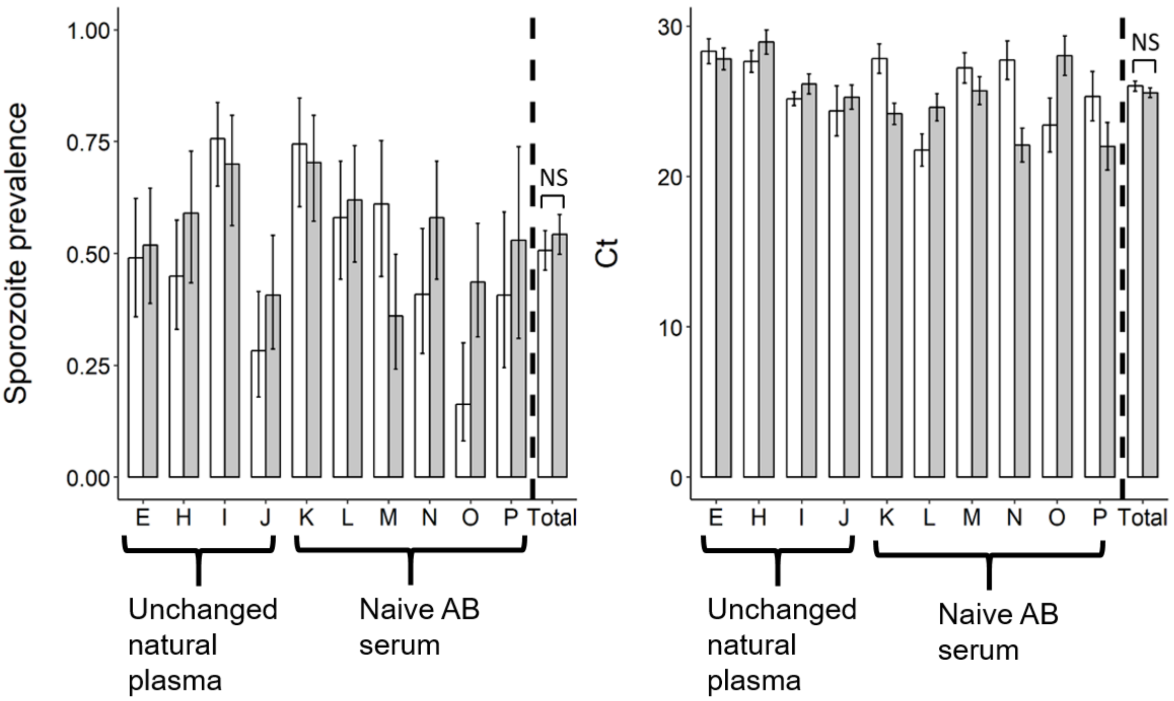


**Figure S2:** Effect of irradiation on sporozoite prevalence (± 95 % CI) at 14 dpbm (left panel), expressed as the number of mosquito head/thoraces detected positive to *Plasmodium falciparum* using qPCR out of the total number of dissected head/thoraces, for each treatment and for 10 parasite isolates; and on sporozoite intensity at 14 dpbm (right panel), expressed as the mean number (± se) of amplification cycle during qPCR (the lower the Ct, the higher the sporozoite intensity) for each treatment and for 10 parasite isolates. * denotes statistically significant difference (P value: 0.01< * < 0.05); NS: not significant.
